# Gut microbiome features are associated with sepsis onset and outcomes

**DOI:** 10.1101/2021.01.08.426011

**Authors:** Krishna Rao, Alieysa R. Patel, Anna M. Seekatz, Christine M. Bassis, Yuang Sun, Oryan Henig, Owen R Albin, John SantaLucia, Robert Woods, Michael A. Bachman

**Affiliations:** Division of Infectious Diseases, Department of Internal Medicine, University of Michigan, Ann Arbor, USA; Department of Pathology, University of Michigan, Ann Arbor, USA; DNA Software, Ann Arbor, USA

**Author notes:** Corresponding author contact information: Michael A. Bachman MD PhD. 7510E MSRB 1, 1301 Catherine St. Department of Pathology. University of Michigan. Ann Arbor, MI 48103.

**Keywords:** Sepsis, microbiota, 23S rRNA gene, 16S rRNA gene, absolute bacterial abundance, qPCR

## Abstract

**Background:** Epidemiologic studies have linked antibiotic exposure to subsequent sepsis, suggesting that microbiome disruption may be in the causal pathway and an independent risk factor. This study tests whether variation in the gut microbiota associates with risk of sepsis onset and its outcomes.

**Methods:** Using a validated surveillance definition, patients with an archived rectal swab from intensive care and hematology units were screened for sepsis. After confirmation by chart review, cases were matched to controls in a 1:2 ratio based on age, gender, and collection date. Relative taxon abundance was measured by sequence analysis of 16S rRNA gene amplicons; total bacterial abundance was measured by qPCR of the 23S rRNA gene. Conditional logistic regression identified clinical and microbiota variables associated with sepsis.

**Results:** There were 103 sepsis cases matched to 206 controls. In a final model adjusting for exposure to broad-spectrum antibiotics and indwelling vascular catheters, high relative abundance (RA) of *Enterococcus* (Odds Ratio (OR) 1.36 per 10% increase, *P*=.016) and high total bacterial abundance (OR 1.50 per 10-fold increase in 23S copies/μL, *P* =.001) were independently associated with sepsis. Decreased RA of butyrate-producing bacteria also independently associated with sepsis (OR 1.20 for 10% decrease in RA, *P* =.041), and mortality in unadjusted analysis (OR=1.47 for 10% decrease in RA, *P*=.034).

**Conclusions:** This study indicates that the microbiota is altered at sepsis onset. The decreased RA of butyrate-producing bacteria in sepsis also associates with mortality, suggesting a therapeutic role for prebiotics and probiotics in the prevention and treatment of sepsis.

**Importance:** Early detection of patients at risk for sepsis could enable interventions to prevent or rapidly treat this life-threatening condition. Prior antibiotic treatment is associated with sepsis, suggesting that disruption of the bacterial population in the gut (the intestinal microbiome) could be an important step leading to disease. To investigate this theory, we matched hospitalized patients with and without sepsis and characterized the patients’ microbiomes close to or at onset of sepsis. We found that several microbiome alterations, including having more total bacteria in the gut was associated with onset, regardless of prior antibiotic treatment. This signature of microbiome disruption brings us closer to identifying the biological causes of sepsis and could be used to develop new diagnostic tests to identify patients at risk of sepsis.

## Introduction

Sepsis, defined as life-threatening organ dysfunction caused by a dysregulated host response to infection, affects 1.7 million people annually in the United States (1, 2). As successfully advocated by the Surviving Sepsis Campaign, to prevent death from sepsis, intervention must be rapid and include a combination of supportive care and treatment of the underlying infection (3). Though advances in care have reduced mortality from 46.9% in the early 1990s to 29% by 2009 (4), this is still an unacceptably high risk of death. Furthermore, prevention efforts have been less successful, as there has been little progress made on overall sepsis incidence, which for example remained stable from 2009–2014 (3) and the worldwide incidence may be significantly higher than previously estimated (5). Prior antibiotic exposure is associated with subsequent sepsis. In particular, antibiotics most disruptive of the gut microbiome as measured by the strength of association with *Clostridioides difficile* infection are high-risk antibiotics for subsequent sepsis (6). This indicates that alterations in the microbiome may be an important sepsis risk factor and a possible target for better preventative and therapeutic efforts

The microbiome may affect sepsis risk by multiple, potentially overlapping, mechanisms. Bacteremia from *Enterococcus* and Proteobacteria are associated with intestinal domination by these taxa (7, 8), suggesting a direct causative link between the microbiome and infections. Alternatively, depletion of butyrate-producing taxa are associated with viral respiratory infections, suggesting the microbiome may also play a more indirect immunomodulatory role (9). Most studies of the microbiome measure relative abundance of bacterial taxa, but their absolute abundance individually and as a community are also associated with various disease states (10) (11). Although there is strong evidence that antibiotics that disrupt the microbiome increase sepsis risk, and that dominance of certain microbiome taxa increase the risk of corresponding infections, the specific pattern of microbiome disruption associated with sepsis onset is unclear. To identify the microbiome variables associated with sepsis and its outcomes, a single-center case-control study was performed using rectal swabs collected before or at the time of sepsis onset.

## Results

### Identifying important potential clinical confounders

#### Cohort characteristics and unadjusted analysis

Using a Centers for Disease Control (CDC) surveillance definition (1) with confirmation by manual chart review, 103 cases and 206 controls were identified among intensive care and hematology oncology patients with an available rectal swab between January 2016–February 2017. Of the sepsis cases, 86 were community onset and 17 were hospital onset, and 73 cases had the rectal swab collected on the same day as starting antibiotics for treatment while 30 had the swab collected in the week prior to starting antibiotics. Among controls, 22 had the swab collected on the same day as, and 6 had the swab collected in the week prior to, starting antibiotics. The remaining 178 controls either did not start antibiotics (66) or started prior to swab collection (112). Cases and controls were well-matched with respect to demographics (**Table 1)**. However, cases had worse baseline vital signs and laboratory values compared to controls, consistent with sepsis, and a higher Elixhauser comorbidity score (12). More cases were exposed to a high-risk antibiotic (third- or fourth-generation cephalosporins, lincosamides, β-lactam/β-lactamase inhibitor combinations, oral vancomycin, and carbapenems) in the 90 days prior to admission.

**Table 1.** Selected baseline characteristics of the study population.

| **Table 1. Selected baseline characteristics of the study population.** |  |  |  |  |
| --- | --- | --- | --- | --- |
| **Variable** | **Cases (n=103)** | **Controls (n=206)** | **OR** | **P** |
| <u>Demographics</u> |  |  |  |  |
| Age (Years) | 59.32±16.79 | 59.18±16.53 | 1.02 | .701 |
| Male | 58(56.31) | 116(56.31) | 1.00 | >.99 |
| White | 86(83.50) | 169(82.04) | 1.62 | .260 |
| Non-Hispanic or Latinx | 98(95.15) | 198(96.12) | 2.50 | .403 |
| <u>Vitals and laboratory values</u> |  |  |  |  |
| Baseline body mass index (kg/m<sup>2</sup>) | 27.98± 6.59 | 30.19± 7.41 | 0.96 | .019 |
| Baseline glucose level (mg/dL) | 196.84 ±145.41 | 156.75± 63.50 | 1.01 | .003 |
| Baseline hemoglobin level (g/dL) | 9.03± 2.50 | 10.60±2.46 | 0.77 | <.001 |
| Baseline white blood cell count (K/μL) | 17.47± 10.79 | 15.77± 47.66 | 1.00 | .723 |
| Baseline platelet level (K/μL) | 169.50± 98.55 | 185.06± 90.74 | 1.00 | .170 |
| Baseline albumin level (g/dL) | 2.89± 0.64 | 3.44± 0.62 | 0.25 | <.001 |
| Baseline albumin > 3.0 (g/dL) | 38 (36.89) | 118 (57.28) | 0.20 | <.00<br>1 |
| Baseline creatinine level (mg/dL) | 2.06± 1.97 | 1.22± 1.25 | 1.50 | <.00<br>1 |
| Baseline heart rate (Beats/min) | 116.52± 23.93 | 100.76±21.37 | 1.03 | <.00<br>1 |
| Baseline systolic blood pressure(mmHg) | 112.41± 23.60 | 119.42 ±19.69 | 0.98 | .010 |
| Baseline diastolic blood pressure (mmHg) | 60.52± 14.23 | 63.85±11.49 | 0.98 | .042 |
| Baseline temperature (F) | 100.16± 1.48 | 99.10± 0.86 | 2.28 | <.00<br>1 |
| Baseline respiratory rate (breaths/min) | 38.55± 14.67 | 25.93± 11.27 | 1.08 | <.00<br>1 |
| Baseline SPO2 (peripheral oxygen saturation) | 95.37± 2.97 | 95.01± 3.68 | 1.04 | .345 |
| Baseline PTT (partial thromboplastin time in seconds) | 39.52± 26.46 | 29.86± 14.23 | 1.02 | .009 |
| Baseline INR | 1.58± 0.96 | 1.20± 0.53 | 2.10 | <.00<br>1 |

| <u>Medications</u> |  |  |  |  |
| --- | --- | --- | --- | --- |
| Total prior antibiotic duration (days) | 13.98 ±19.32 | 14.37± 23.27 | 1.00 | .943 |
| Total number prior concurrent antibiotic classes | 2.31± 1.40 | 1.86± 1.43 | 1.10 | .667 |
| High-risk antibiotic exposure at baseline | 27 (26.21) | 19 (9.22) | 3.43 | <.001 |
| Proton pump inhibitor | 53 (51.46) | 82 (39.81) | 1.61 | .053 |
| Probiotics | 3 (2.91) | 1 (0.49) | 6.00 | .121 |
| Immunosuppressant | 34 (33.01) | 45 (21.84) | 1.77 | .036 |
| Immunomodulator | 8 (7.77) | 16 (7.77) | 1.00 | >.99 |
| H2 receptor blocker | 52 (50.49) | 73 (35.44) | 1.85 | .013 |
| Antipsychotic | 26 (25.24) | 37 (17.96) | 1.52 | .144 |
| <u>Comorbidities</u> |  |  |  |  |
| Cardiac arrhythmias | 67 (65.05) | 76(36.89) | 3.20 | <.001 |
| Chronic pulmonary disease | 38 (36.89) | 40 (19.42) | 2.39 | .001 |
| Coagulopathy | 31 (30.10) | 35 (16.99) | 1.94 | .014 |
| Congestive heart failure | 39(37.86) | 46 (22.33) | 2.22 | .005 |
| Iron deficiency anemia | 17 (16.50) | 12 (5.83) | 3.14 | .004 |
| Diabetes | 36(34.95) | 46(22.33) | 1.89 | .021 |
| Fluid electrolyte disorders | 78 (75.73) | 55 (26.70) | 10.40 | <.001 |
| Liver disease | 27 (26.21) | 31 (15.05) | 2.06 | .019 |
| Metastatic cancer | 9 (8.74) | 35 (16.99) | 0.48 | .059 |
| Other neurological disorders | 28 (27.18) | 11(5.34) | 5.82 | <.001 |
| Paralysis | 6 (5.83) | 2 (0.97) | 6.00 | .029 |
| Psychoses | 7 (6.80) | 4 (1.94) | 4.22 | .039 |
| Renal failure | 32 (31.07) | 35 (16.99) | 2.26 | .005 |
| Solid tumor without metastasis | 17 (16.50) | 63 (30.58) | 0.45 | .010 |
| Valvular disease | 18 (17.48) | 53 (25.73) | 0.56 | .082 |
| Weight loss | 25 (24.27) | 23 (11.17) | 2.58 | .004 |
| Elixhauser score | 6.82± 2.39 | 4.75± 2.84 | 1.32 | <.001 |
| High_Elixhauser(elixhauser_score >7) | 33(32.04) | 32 (15.53) | 2.46 | .001 |

| <u>Devices</u> |  |  |  |  |
| --- | --- | --- | --- | --- |
| Urinary catheter at baseline | 81 (78.64) | 116 (56.31) | 2.85 | <.00<br>1 |
| Indwelling venous catheter at<br>baseline | 79 (76.70) | 88 (42.72) | 4.33 | <.00<br>1 |
| Feeding tube at baseline | 56 (54.37) | 70 (33.98) | 2.32 | <.00<br>1 |

#### Modeling sepsis with clinical variables

To determine which clinical variables were independently associated with sepsis, a model was derived by backward elimination (**Table 2**). High-risk antibiotic exposure, fluid & electrolyte disorder, and indwelling venous catheter at baseline were independent risk factors for sepsis. There were also inverse associations between peripheral vascular disorder and valvular disease and sepsis that could not be explained within the limits of the dataset and retrospective data collection. The model fit the data well, with an area under the receiver operator curve (AUROC) of 0.93 (**Supplemental Figure 1**). These variables were then considered as potential confounders as we tested hypotheses about microbiota features associated with sepsis.

**Table 2.** Final clinical model for sepsis.

| **Table 2. Final clinical model for sepsis.** |  |  |
| --- | --- | --- |
| **Variable** | **OR [95%CI]** | ***P*** |
| High-risk antibiotic exposure at baseline | 4.10 [1.52, 11.07] | .005 |
| Fluid & Electrolyte Disorder | 10.86 [4.23, 27.92] | <.001 |
| Other Neurological Disorders | 6.52 [2.12, 19.99] | .001 |
| Peripheral Vascular Disorder | 0.22 [0.07, 0.68] | .009 |
| Cardiac Arrhythmias | 2.74 [1.21, 6.21] | .016 |
| Valvular Disease | 0.24 [0.09, 0.68] | 0.007 |
| Indwelling venous catheter at baseline | 5.78 [2.41, 13.85] | <.001 |

### Identifying microbiota features associated with sepsis

#### Community structure, and sepsis

We measured Shannon diversity and observed a threshold effect, defining “low Shannon diversity” at an optimal cut point of <2.5. Low Shannon diversity associated significantly with sepsis on unadjusted analysis (OR=1.79, *P* =.024, **Supplemental Table 3**).

There was a significant difference in the relative abundances of shared and non-shared OTUs of samples from sepsis cases compared to controls (beta-diversity) as measured by Analysis of Molecular Variance (AMOVA) on θ_YC_ distances (represented by PCoA, **Supplemental Figure 2**; *P* <.001) (13, 14). There was no significant difference between the microbiota of samples taken from sepsis patients on the day of sepsis diagnosis (n=73) versus the week prior to sepsis diagnosis (n=30) (*P =*.904) and both were significantly different than controls (*P* <0.001 and *P* =.004, respectively). There was also no difference between the microbiota of community onset (n=86) and hospital onset cases (n=17) by AMOVA (*P =* 0.566) and both were significantly different than controls (*P* <.001 and *P =*.005, respectively). Individual subject scores from PCoA axes 1 and 3 were significantly associated with sepsis (**Supplemental Table 3**). Using Partitioning Against Medoids (PAM) clustering based on Jensen-Shannon Divergence and without regard to case status, the samples clustered into 2 community types (optimal partitioning based on highest Laplace value, testing up to 5 partitions), and with community type 2 associated with sepsis (**Supplemental Table 3**) (15, 16).

#### High-rank taxonomic and constructed variables vs. sepsis

High relative abundance of the Bacteroidetes phylum and the *Enterobacteriaceae* family associated with sepsis, while relative abundance of the Firmicutes was higher in controls. Neither the microbiome health index (relative abundance of [Bacteroidia + Clostridia]/[ γ-Proteobacteria + Bacilli]) (17) nor the Firmicutes/Bacteroidetes ratio associated with sepsis (**Supplemental Table 3**). However, the relative abundance of butyrate-producing bacteria (**Supplemental Dataset**; (9) was inversely associated with sepsis (OR=0.77 for every 10% increase, *P=*.001).

#### Individual bacterial taxa and sepsis

Linear discriminant analysis Effect Size (LEfSe) (18) analysis of the individual OTUs enriched in the two different community types and in sepsis cases vs. controls revealed that OTU #2 / *Enterococcus* was enriched in both sepsis cases and microbiota community type 2 (**Supplementary Results, Supplemental Figure 3)**. Although LEfSe does not account for matching between cases and controls, it is a robust method that avoids type I statistical errors without a significant reduction in statistical power. As a complementary approach, unadjusted analyses of OTUs using conditional logistic regression accounting for case/control matching was performed (**Supplemental Table 3).**The presence and abundance of specific OTUs, in particular relative abundance of OTU #2 / *Enterococcus*, were associated with higher odds of sepsis, though these analyses did not reach statistical significance.

#### Total bacterial abundance

To measure total bacterial abundance in rectal swab samples, we developed a PCR assay for 23S rRNA by compiling a focused list of the organisms most prevalent in stool and using *PanelPlex* and *ThermoBLAST* software (DNA Software, Inc., Ann Arbor, MI) to find optimal consensus primers and probes (**Supplemental Results, Supplemental Table 1**) (19, 20). This assay demonstrated a significant association on unadjusted analyses between increased total bacterial abundance and sepsis (OR 1.67 for every 10-fold increase in 23S gene copies/ μL, *P* <.001, **Supplemental Table 3**).

### Holistic modeling of sepsis risk with both clinical and microbiome variables

Analysis of the clinical variables for confounding (associated both with sepsis and one or more microbiota variables) identified multiple co-morbidities and exposure to high risk antibiotics as potential confounders to include in our adjusted models (**Supplemental Table 4)**. To avoid an unstable and overfit model caused by too many candidate variables, high Elixhauser score was carried forward as a measure of comorbidity burden into the models instead of individual comorbidities. Since it was not selected for inclusion, we forced “high Elixhauser score” back into this model to control for comorbid disease, and it neither changed the results of the other covariates nor was significant itself (*P* =.771), so it was not included in the final model containing two clinical and two microbiota variables (**Table 3)**. After adjusting for indwelling vascular catheter and prior high-risk antibiotic exposure, we found that for every 10-fold increase in 23S rRNA gene copies/μL, there was a 1.50-fold increased odds of sepsis. That is, the odds of sepsis rose as bacterial abundance rose. Additionally, every 10% increase in *Enterococcus* relative abundance results in a 1.36-fold increased odds of sepsis. Thus, these microbiota-derived variables were the best independent predictors of sepsis when considered alongside clinical predictors. Given its functional and potential therapeutic significance, we separately tested the hypothesis that decreased relative abundance of butyrate-producing bacteria was associated with sepsis, and found it was associated with sepsis independent of indwelling vascular catheter and high risk antibiotic exposure (OR 1.2 for every 10% decrease; **Supplemental Table 5**) (9).

**Table 3.** Final multivariable model for sepsis.

| **Table 3. Final multivariable model for sepsis.** |  |  |
| --- | --- | --- |
| **Variable** | **OR [95% CI]** | ***P*** |
| Indwelling vascular catheter | 4.75 [2.5, 9.0] | <.001 |
| High risk antibiotic exposure at baseline | 3.52 [1.5, 8.1] | .003 |
| 23S rRNA gene copies/μL (per 10-fold increase) | 1.50 [1.2, 1.9] | .001 |
| OTU #2 / *Enterococcus* (for every 10% increase in abundance) | 1.36 [1.1, 1.8] | .016 |

### Identification of microbiota features associated with outcomes following sepsis

#### Mortality among sepsis patients and microbiota factors

Among the 103 subjects with sepsis, 28 (27.2%) died. In an exploratory analysis of predictors of mortality, gut microbial community type 2 (OR=5.40, *P*=.03) and decreased relative abundance of butyrate-producing bacteria (OR=1.47 per 10%, P=0.034) were strongly associated with mortality among septic patients. In contrast, scores from PCoA axis 1 and increased relative abundance of *Peptoniphilus* species had protective effects on unadjusted analyses **(Supplemental Table 6).**Modeling, limited by sample size, only selected 2 variables, and after accounting for age, having a gut community type 2 retained borderline statistical significance for increased mortality risk (OR=4.48, *P=*.057, **Supplemental Table 7**). Adding relative abundance of butyrate-producing bacteria into this model, it had borderline significance as a protective factor **(Supplemental Table 8)**.

## Discussion

Our finding that several features of the gut microbial community are independently associated with sepsis supports the hypothesis that the gut microbiome, at least in part, mediates sepsis risk (6). Our most notable findings are that within a week of sepsis onset, higher total bacterial abundance, higher relative abundance of OTU #2 (*Enterococcus* species), and lower relative abundance of butyrate-producing bacteria all associated with increased odds of sepsis. These findings held even after adjustment for potential clinical confounders, including exposure to high-risk antibiotics associated with sepsis (6).

The association with increased total bacterial abundance, as measured by 23S rRNA gene qPCR, is particularly intriguing. Since 16S rRNA gene sequencing alone only allows for calculation of relative abundance, absolute quantification of taxa is less commonly reported. Based on measuring DNA concentration normalized to total weight of fecal samples, patients with recurrent *C. difficile* infection had lower overall bacterial density compared to those with non-recurrent disease, and this lower overall bacterial density was restored by fecal transplant (10). Our findings suggest that sepsis may be associated with higher absolute bacterial abundance. Alternatively, there may be differences in the amount of fecal material in the rectal and perirectal area of patients that develop sepsis, which could be attributed to physiological variables (e.g. anal sphincter function) or hygiene. There is also likely variation in sample collection, which could lead to differences in measured total bacterial abundance. However, this variation is likely to be random, and we have also observed an association between *Enterobacterales* total abundance and infection (11). Thus, our findings should be confirmed in a study that utilizes stool samples.

The biological reason for the association between *Enterococcus* abundance and sepsis is unclear. Previous studies have shown that *Enterococcus* dominance is associated with subsequent vancomycin-resistant *Enterococcus* bacteremia (7). Furthermore, there is evidence that dominance by *Enterococcus* may be associated with poor patient health (21), and with loss of colonization resistance to resistant Gram-negative pathogens (22).

We also found that a lower relative abundance of butyrate-producing bacteria was associated with sepsis. Butyrate has been associated with immunomodulatory effects in the intestine and on lung infections (9), and could have similar protective effects against sepsis. This separate finding hints at functional disruptions of the microbiome that could be therapeutic targets for reduction of sepsis using probiotics and/or prebiotics as demonstrated for prevention of neonatal sepsis (23).

The clinical model (**Table 2**) confirmed that high-risk antibiotics were associated with sepsis, as observed in the epidemiologic study by Baggs et al (6). We also found a positive association with fluid and electrolyte disorders and neurologic disorders, which we previously found to be associated with *Klebsiella pneumoniae* infections from this same patient population (24). The reason for the inverse association between sepsis and both peripheral vascular disorder and valvular disease was unclear. However, we also observed this inverse correlation between peripheral vascular disorder and *K. pneumoniae* infection previously (25). These negative associations could indicate that patients with these disorders are admitted for surgery and are a distinct subset of the ICU population at lower risk of infections, and this deserves further study.

Our study has several limitations. Though a major problem nationwide, at the level of our individual health system, sepsis is still a rare outcome. This necessitated a case-control design in lieu of a more methodologically straightforward cohort study, and there are inherent concerns regarding information bias, confounding, and data reliability in any retrospective study. We attempted to mitigate these limitations through manual chart review, matching, and careful, adjusted modeling. Ideally all of our rectal swab samples would have been collected before sepsis onset and before any antibiotics were started, but we were limited in the samples available to us and many were obtained on the day of sepsis onset. It is reassuring that there were no substantial differences in overall community structure, as measured by beta-diversity, when comparing samples from the week prior to sepsis to ones obtained on the day of onset (*P* =.904). Although we identified the microbiome-derived variables most strongly associated with sepsis after adjustment for clinical confounders, some of the other microbiota associated variables we identified on unadjusted analysis may also be important (**Supplemental Table 3**), and there may be others of importance that we did not have sufficient power to detect at all. This may be due to confounding from clinical variables or lack of sufficient power for model inclusion in the setting of other, more explanatory variables, which could be addressed in a future study with a larger sample size.

In conclusion, our study is consistent with the hypothesis that the gut microbiome in part mediates risk of sepsis and its subsequent outcomes. These findings also have immediate feasibility for monitoring and prediction, as the final model incorporates information that is easily obtainable during a patient’s hospitalization and we currently have a qPCR design that measures total rectal bacterial abundance that is associated with sepsis. This could be paired with an *Enterococcu*s-specific qPCR to measure relative abundance and obtain the 4^th^ variable in the model. Such detection of high-risk patients, if achieved rapidly and cheaply, can enable trials of infection prevention interventions.

## Methods

### Study design

This was a nested case-control study within a retrospective cohort of intensive care and hematology/oncology patients with archived rectal swabs. Based on a power calculation, the enrollment goal was 100 cases of sepsis matched to 200 controls (see **Supplementary Methods**). Electronic medical record data from patients with an archived rectal swab sample obtained between January 2016 and February 2017 were screened by Sepsis-3 criteria (1) (2). Sepsis was defined as 1) presumed serious infection indicated by obtaining blood cultures; 2) 4 days of antibiotic treatment started ±2 days from blood cultures; and 3) acute organ dysfunction present ±2 days from blood cultures (1). Each case screening positive was confirmed with manual review by an infectious diseases attending physician (KR). Controls were excluded if they had evidence of infection or an ICD-9 code for sepsis that did not meet criteria upon manual review. Cases were matched to eligible controls based on age (+/− 10 years), sex, and date of swab collection (+/− 45 days).

For all cases and controls, detailed electronic medical record data was extracted. Comorbidities and Elixhauser scores were extracted and calculated as previously described (12). Baseline values for labs and vitals were defined as the either the maximum (e.g. temperature) or minimum value (e.g. albumin) within 48 hours of admission. Antibiotic exposure metrics for the 90 days prior to admission included total duration, the number of concurrent classes of antibiotics, and risk category, defined as high, medium, or low based on prior association with both microbiome disruption and sepsis (6). High-risk antibiotics were defined as third and fourth-generation cephalosporins, lincosamides, β-lactam/lactamase inhibitors, oral vancomycin, carbapenem, daptomycin, and metronidazole (the latter changed from low in the original Baggs *et al.* study, (6)based on personal communication with the authors). Medium risk antibiotics included penicillins, aminoglycosides, and intravenous vancomycin. Low risk antibiotics were first and second-generation cephalosporins, macrolides, tetracyclines, sulfa antibiotics, and fluoroquinolones (the latter changed from high in the original Baggs *et al*. study, based on personal communication with the authors).

### Bacterial community analysis

Rectal swab analysis was performed on 250 μL of liquid Amies transport media in the E-swab transport system (Becton Dickinson, Franklin Lakes, NJ). Total DNA extraction, library preparation, and 16S rRNA gene-based sequencing using Illumina technology were conducted by the University of Michigan Microbial Systems Laboratory (MSL) using dual-indexing sequencing strategy targeting the 16S rRNA V4 region(26). The resulting sequences were processed and analyzed using mothur v1.39.5 (www.mothur.org/wiki/MiSeq_SOP) (26, 27). Analysis was performed on 309 samples from 103 complete strata with a minimum of 2500 sequences per sample. Bacterial abundance was measured by 23S rRNA gene qPCR on an aliquot of the same DNA used for sequencing. Details are described in **Supplemental Methods**.

### Modeling

The primary outcome was sepsis and the primary predictors of interest were features of the gut microbiota. We assessed for threshold effects and, where present, reconstructed the variables (e.g. dichotomization). In addition to diversity and richness, we considered various taxonomic variables such as phylum (focus on Bacteroidetes, Firmicutes) and class (focus on Bacilli, Clostridia, Bacteroidia, and γ-Proteobacteria). The OTUs were modeled both as relative abundance and as absent/present. To reduce false positives, we focused only on the filtered list of OTUs and applied a false discovery rate correction. Only OTUs with a corrected *P* value <.2 were carried forward for consideration in adjusted models. Conditional logistic regression was used for both the unadjusted and adjusted analyses to test our hypotheses. To build models, we only included clinical covariates that were flagged as potential confounders (i.e. associated *both* with the microbial predictor and sepsis) and performed backward elimination with a likelihood ratio test (cutoff of *P* <.05 for retention). We assessed for interactions in the final models and included them if statistically significant.

**Figure 1.**
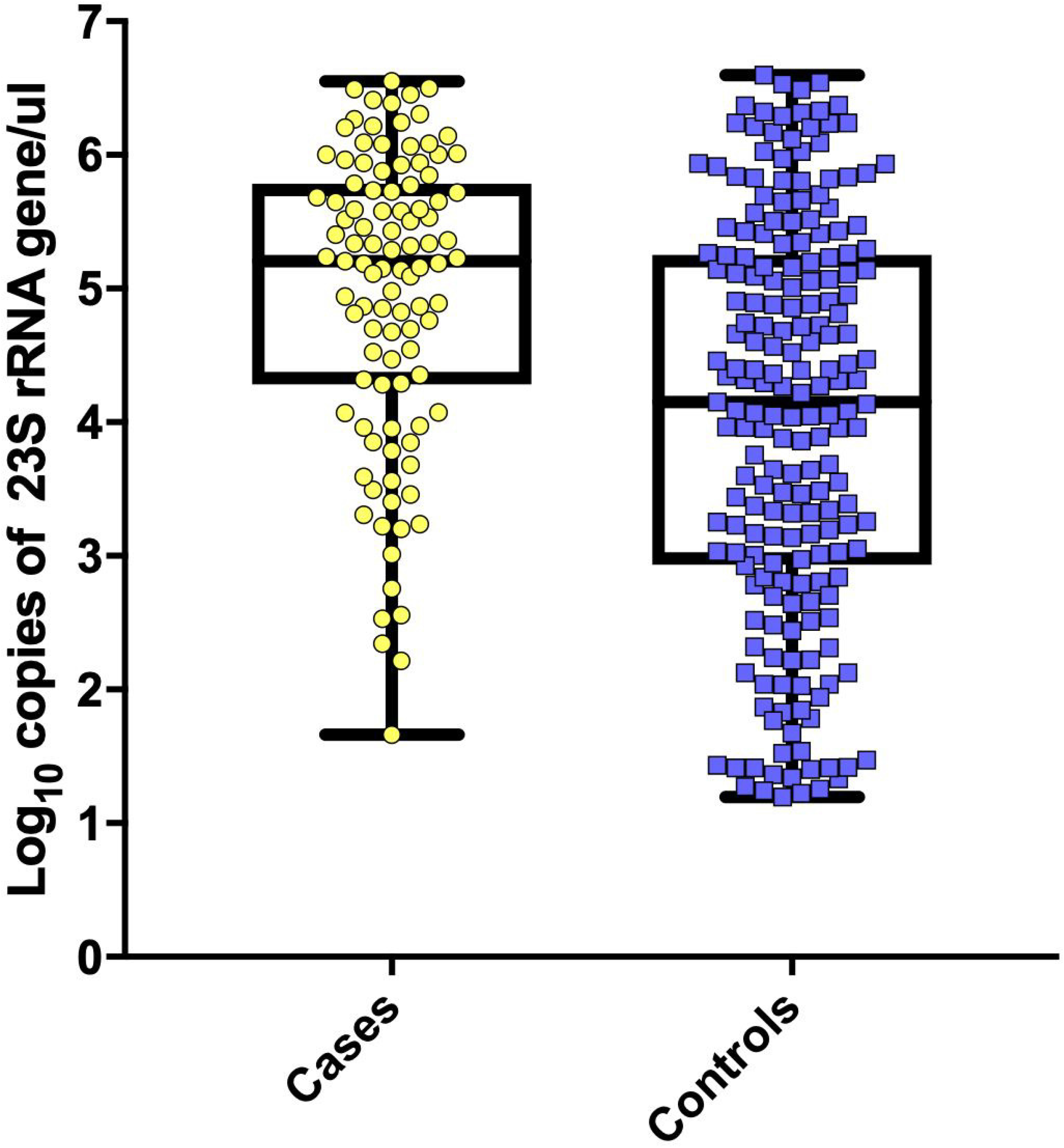
Increased total bacterial abundance, measured by 23S rRNA gene qPCR, is associated with sepsis. 23S rRNA gene from rectal swabs of cases of sepsis (n=103) and matched controls (n=106) was amplified by qPCR and quantified in gene copies/ul relative to a standard curve of *Klebsiella pneumoniae* KPPR1 genomic DNA. Box and whiskers plots showing median, interquartile ranges, minimum and maximum values are shown. *P* <0.001 in unadjusted logit analysis.

## Acknowledgements

We would like to thank the University of Michigan Data Office for assistance with electronic medical record queries.

## Data Availability

All sequencing data has been deposited in the Sequence Read Archive. See Supplemental Dataset 2 for accession numbers and associated metadata.

## Supplemental Materials

**Supplemental Results and Methods.** Additional results and methodological details.

**Supplemental Dataset 1.** Sequences and taxa used for PCR design and identification of butyrate producing taxa.

**Supplemental Dataset 2.** Sequence Read Archive accession numbers and associated metadata.

